# 1LocusSim a mobile-friendly simulator for teaching population genetics

**DOI:** 10.1101/2023.06.20.545745

**Authors:** Antonio Carvajal-Rodríguez

## Abstract

**Summary:** Biology students often struggle with the fundamental concepts of evolutionary genetics, including genetic drift, mutation, and selection. To address this problem, 1LocusSim was developed to simulate the interaction of different factors, such as population size, mutation, selection, and dominance, to study their effect on allelic frequency during evolution. With 1LocusSim, students can compare theoretical results with simulation outputs and solve and analyze different problems of population genetics. The 1LocusSim web has a responsive design which means that it has been specifically designed to be used on smartphones. To demonstrate its use, I review the classical overdominance model of population genetics and highlight a characteristic that is often not explicitly stated. Specifically, it is emphasized that the equilibrium of the model does not depend on the homozygous selection coefficients but rather on the ratio of the selection coefficients. This is already clear from the classical formula but maybe not so much for students. Also it implies that the equilibrium can be expressed solely in terms of the dominance coefficient *h*. To verify these theoretical prediction, I utilize the simulator and calculate the equilibrium for the well-known case of sickle cell anaemia.

Simulating basic population genetic models on smartphones can be a powerful learning aid that fosters critical skills and opens up new opportunities for Biology students. By utilizing this tool, students can learn at their own pace and convenience, anywhere and anytime.

**Software and data availability:** 1LocusSim if freely available at https://1LocusSim-biosdev.pythonanywhere.com/. Website implemented under the Bottle micro web-framework for Python, with all major browsers supported.

**Supplementary information:** The manual and examples are available from the help button of the program or directly at https://acraaj.webs.uvigo.es/1LocusSim/1LocusSim_EN.html. Both the program and the manual pages are in English and Spanish.

## Introduction

Smartphones offer new learning opportunities and provide approaches to foster the development of critical skills for learners in the 21st century. Smartphones have become an ingrained aspect of students’ lives, making it a comfortable medium for learning. One of the key benefits of the smartphone is its portability, allowing students to access learning content anytime, anywhere. With learning available at their fingertips, students can expand their knowledge without interruption. The prevalence of smartphones in households of all demographics makes it a readily available and accessible tool for learning. Here, I present a one-locus population genetics simulator called 1LocusSim. Unlike other teaching software, 1LocusSim is a wep app so it is not limited to a specific platform or operating system, as it is available in the cloud and does not require installation. It can be accessed from any device, including smartphones, for which it has been specially designed, and comes with a set of companion web pages that include a glossary of basic concepts, formulas, examples, and exercises to enhance the students’ learning experience. Therefore, the student only needs to type the web address (1LocusSim-biosdev.pythonanywhere.com) from their preferred device to run the program.

As an example of the use of the simulator, we will review the basics of the overdominance model to calculate the equilibrium value of the sickle cell gene and use 1LocusSim to perform simulations that confirm the theoretical result.

### Two allele overdominance model

In an overdominance fitness model, the heterozygous has higher fitness than the corresponding homozygotes. Typically, this is represented as the heterozygous having fitness 1, while the A_1_A_1_ and A_2_A_2_ homozygotes have lower fitness given by 1-*s*_1_ and 1-*s*_2_, respectively. In a large population, the equilibrium value for that model does not require mutation (Crow and Kimura, 1970) and it depends only on the ratio *k* = *s*_2_ /*s*_1_ of the selection coefficients (see equation S1 in the Supplementary Appendix and the [overdominance](https://acraaj.webs.uvigo.es/1LocusSim/1LocusSim_Seleccion_EN.html) section of the online manual). Sometimes it’s useful to express the overdominance model using the selection coefficient *s* and the dominance coefficient *h* from the classical selection model, as shown in Table 1 (and in [Manual](https://acraaj.webs.uvigo.es/1LocusSim/1LocusSim_Seleccion_EN.html) Tables 2 and 3).

**Table 1.**
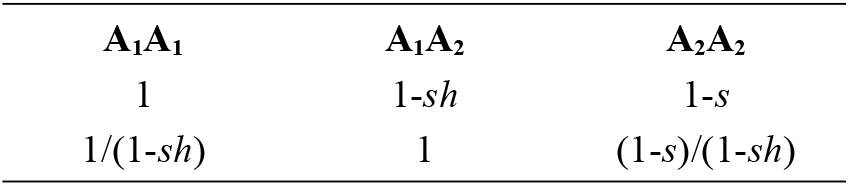
Overdominance expressed from the model with selection coefficient *s* and dominance coefficient *h* (*s*>0, *h*<0 or *s*<0, *h*>1)

**Table 2.**
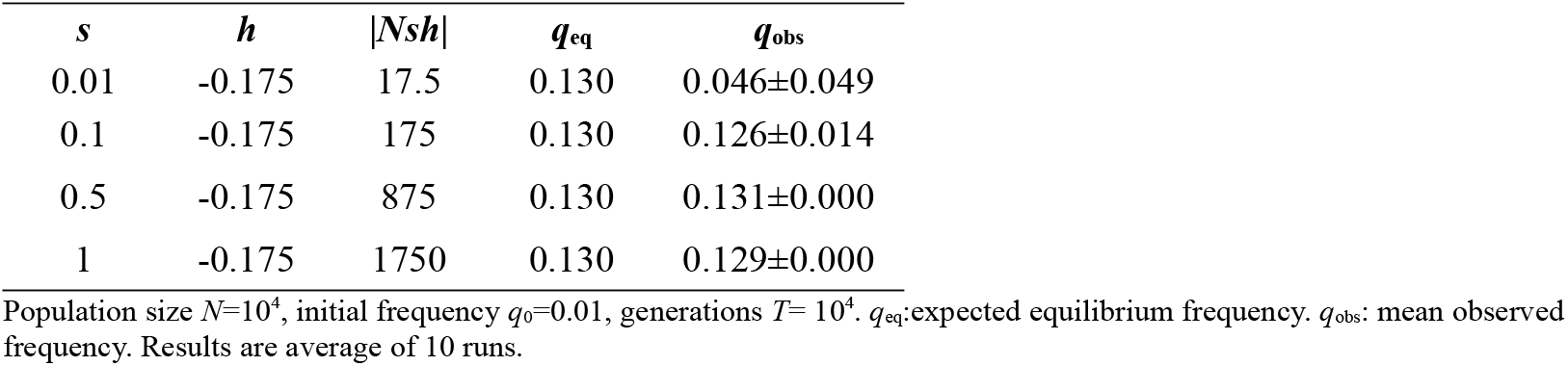
Equilibrium values for the sickle cell gene after simulation

| $s$ | $h$ | $\vert Nsh\vert $ | $q_{eq}$ | $q_{obs}$ |
| --- | --- | --- | --- | --- |
| 0.01 | -0.175 | 17.5 | 0.130 | 0.046±0.049 |
| 0.1 | -0.175 | 175 | 0.130 | 0.126±0.014 |
| 0.5 | -0.175 | 875 | 0.130 | 0.131±0.000 |
| 1 | -0.175 | 1750 | 0.130 | 0.129±0.000 |

Population size $N=10^4$ , initial frequency $q_0=0.01$ , generations $T= 10^4$ . $q_{eq}$ : expected equilibrium frequency. $q_{obs}$ : mean observed frequency. Results are average of 10 runs.

In this case, *h* = 1/(1-*k*) and the equilibrium value (supplementary equation S2) can be expressed just in terms of the dominance coefficient (Caballero, 2020). For finite population sizes (Robertson, 1962) showed that the effect of genetic drift is controlled by *N*(s_1_+s_2_)*f* where *f* is the equilibrium frequency of the lowest fit allele. Under our notation the condition in Robertson can be expressed as a function of |*Nsh*|. The higher this value, the lower the effect of genetic drift and *vice versa*.

### Simulations

Since 1LocusSim is designed to be used both in classrooms and on mobile devices such as smartphones and tablets, it is important that simulation times don’t exceed a couple of minutes even in the worst-case scenario. Therefore, the parameter values have been limited to ensure that the simulation runs within this time frame (please refer to the online manual for a more detailed explanation of parameter limits). Despite these limitations, we can use 1LocusSim to test the dependence of overdominance equilibrium solely on the *h* parameter and study as an example the Sickle cell anaemia overdominance model.

Therefore, we first test whether the simulations behave as predicted for the overdominance model in general, such that the equilibrium values only depend on *h*. The simulation results shown in supplementary Table S1, demonstrate that the equilibrium values are in good agreement with expectations and clearly depend on the *h* value as predicted (see also Manual’s Table 4 and Figure 6). However, as expected, the lower the |*Nsh*| value the higher the noise due to genetic drift.

### Overdominance equilibrium for a lethal allele: The sickle cell anaemia

Sickle-cell anaemia (SCA) is a classical example of overdominance. In high-malaria regions of Africa, homozygotes for the non-sickle-cell anaemia allele have a relative fitness of 0.85 (*s*_1_=0.15). The SCA allele can be considered a recessive lethal (*s*_2_=1) in regions without modern medical care (Hartl, 2020). Heterozygous individuals have fewer physiological effects of the SCA condition and partial resistance to malaria. Therefore, the overdominance model for the SCA implies, using the previous notation, *k =s*_2_/*s*_1_*=*6.7 and *h* = 1/(1-*k*) =-0.175 and the expected equilibrium frequency of the SCA allele in the absence of mutation is *q*_eq_= *h*/(2*h-*1) = 0.13.

Now, we can run simulations to calculate the equilibrium value of the sickle cell anaemia gene. Thus, we set the following parameters in LocusSim: *n*=1, *N*=10^4^, *q*_0_=0.01, μ=0, *h*=-0.175, *T*= 10^4^ and run simulations under different selection coefficients (*s*=0.01, 0.1, 0.5 and 1). We repeat each simulation 10 times and compute the mean and standard deviation of the observed frequency *q*_obs_. Simulations confirm that an equilibrium value close to 0.13 is obtained for *h*=-0.175 under different selection coefficients when |*Nsh*| is high enough (Table 2).

In addition, the equilibrium with mutation (Bürger, 1998) was also obtained (equation S3 in Appendix) and the equilibrium predicted values confirmed by simulation (Table S2).

## Discussion

The 1LocusSim simulator usefulness has been demonstrated by reviewing the fundamentals of the overdominance model. Additional examples for mutation, drift and selection equilibriums jointly with exercises, and detailed explanations can be found in the online manual.

Many Biology students often feel a strong rejection towards mathematics and related knowledge. This is often attributed to difficulties in understanding abstract concepts, whether they are mathematical or from other fields, as well as a lack of awareness regarding its relevance in Biology. Moreover, the approach to knowledge acquisition in the field of Biology tends to have limited emphasis on quantitative aspects (Chiel *et al*., 2010; Feser *et al*., 2013; Rubinstein and Chor, 2014). The utilization of smartphones for simulating fundamental population genetic models can prove to be an effective educational tool, promoting critical skills and unlocking novel prospects for Biology students. 1LocusSim enables students to learn at their preferred pace and convenience, anywhere and anytime.

## Acknowledgements

I thank A. Caballero and E. Rolán-Alvarez and two anonymous reviewers for their helpful comments on the program and the manuscript.

## Appendix

### Two allele overdominance model

In an overdominance fitness model, the heterozygote has higher fitness than the corresponding homozygotes. Typically, this is represented as the heterozygote having fitness 1, while the A_1_A_1_ and A_2_A_2_ homozygotes have lower fitness given by 1-*s*_1_ and 1-*s*_2_, respectively.

In a large population, the equilibrium value for that model does not require mutation and is (Crow and Kimura, 1970) *q*_eq_ = *s*_1_ / (*s*_1_ +*s*_2_). This equation can be rearranged as

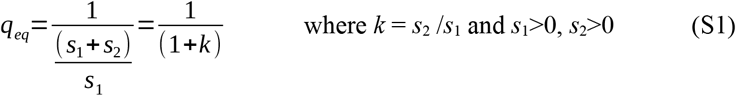

Note that *k* is only defined in an overdominance scenario, i.e. only if *s*_1_ and *s*_2_ > 0. From (S1) it is clear that the equilibrium depends on the ratio of the selection coefficients rather than on each coefficient. Different pairs *s*_1_, *s*_2_ may have the same equilibrium frequency provided the ratio remains constant. The equilibrium have different ranges depending on if *k*>1 ie. *w*_A1A1_ > *w*_A2A2_, *q*_eq_ is in (0, 0.5) on the contrary, if *k<*1 ie. *w*_A1A1_ < *w*_A2A2_, *q*_eq_ is in (0.5, 1). If *k* = 1, *q*_eq_ = 0.5.

The overdominance model can be expressed using the selection coefficient *s* and the dominance coefficient *h* from the classical selection model, as has been shown in Table 1 in the manuscript. The Table 1 presents an overdominance model when *s*>0, *h*<0 or *s*<0, *h*>1. Consequently we can express *k* = *s*_2_/*s*_1_=(*h*-1)/*h* or *h* = 1/(1-*k*) and so the equilibrium value

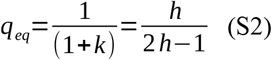

It’s worth noting that the equilibrium value is determined by the dominance coefficient rather than the selection coefficient (Caballero, 2020). This is because the equilibrium depends on the ratio of the homozygous coefficients, rather than the coefficients themselves. For finite population sizes (Robertson, 1962) showed that the effect of genetic drift is controlled by *N*(s_1_+s_2_)*f* where *f* is the equilibrium frequency of the lowest fit allele. Under our notation the condition in Robertson can be expressed as a function of |*Nsh*|. The higher this value, the lower the effect of genetic drift and *vice versa*.

**Table S1.** Overdominance: comparison of expected and observed equilibrium values.

| Model | $s$ | $h$ | $\vert Nsh\vert $ | $q_{eq}$ | $q_{obs}$ |
| --- | --- | --- | --- | --- | --- |
| $w_{A1A1} > w_{A2A2}$ | 0.01 | -0.1 | 10 | 0.083 | 0.021±0.198 |
| $w_{A1A1} > w_{A2A2}$ | 0.1 | -0.1 | $10^2$ | 0.083 | 0.074±0.028 |
| $w_{A1A1} > w_{A2A2}$ | 1 | -0.1 | $10^3$ | 0.083 | 0.086±0.000 |
| $w_{A1A1} > w_{A2A2}$ | 0.01 | -0.5 | 50 | 0.25 | 0.272±0.036 |
| $w_{A1A1} > w_{A2A2}$ | 0.1 | -0.5 | 500 | 0.25 | 0.243±0.010 |
| $w_{A1A1} > w_{A2A2}$ | 1 | -0.5 | $5 \times 10^3$ | 0.25 | 0.251±0.000 |
| $w_{A1A1} > w_{A2A2}$ | 0.01 | -100 | $10^4$ | 0.4975 | 0.498±0.000 |
| $w_{A1A1} > w_{A2A2}$ | 0.1 | -100 | $10^5$ | 0.4975 | 0.498±0.000 |
| $w_{A1A1} > w_{A2A2}$ | 1 | -100 | $10^6$ | 0.4975 | 0.496±0.000 |
| $w_{A1A1} < w_{A2A2}$ | -0.01 | 1.05 | $10^2$ | 0.955 | 0.968±0.056 |
| $w_{A1A1} < w_{A2A2}$ | -1 | 1.05 | $10^4$ | 0.955 | 0.954±0.01 |
| $w_{A1A1} < w_{A2A2}$ | -0.01 | 1.25 | $1.25 \times 10^2$ | 0.83 | 0.84±0.049 |
| $w_{A1A1} < w_{A2A2}$ | -1 | 1.25 | $1.25 \times 10^4$ | 0.83 | 0.836±0.01 |
| $w_{A1A1} < w_{A2A2}$ | -0.01 | 100 | $10^4$ | 0.5025 | 0.506±0.000 |
| $W_{A1A1} < W_{A2A2}$ | -1 | 100 | $10^6$ | 0.5025 | 0.503±0.000 |

Population size $N=10^4$ , initial frequency $q_0 = 0.01$ , number of generations $T= 10^4$ . Results are average of 10 runs.

#### Equilibrium with overdominance and mutation

We can use the equation (2.5) in (Bürger, 1998) with *h*<0 to obtain the equilibrium value with overdominance for a lethal allele (*s*=1), mutation μ>0 and negligible retromutation *ν*=0. After some algebra manipulation we obtain

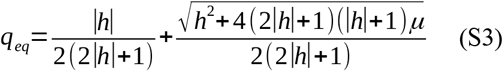

Then if we substitute *h*=-0.175 the equilibrium value is

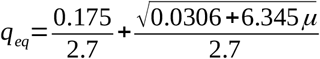

if μ=5×10^−8^ (Melamed *et al*., 2022) then

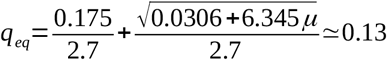

if μ=0.01 then *q*_eq_ = 0.178 and if μ=0.1 then *q*_eq_ = 0.367.

**Table S2.** Sickle cell anaemia equilibrium with mutation.

| $s$ | $h$ | $\mu$ | $q_{eq}$ | $q_{obs}$ |
| --- | --- | --- | --- | --- |
| 1 | -0.175 | $5 \times 10^{-8}$ | 0.130 | 0.130±0.000 |
| 1 | -0.175 | 0.01 | 0.178 | 0.173±0.000 |
| 1 | -0.175 | 0.1 | 0.367 | 0.347±0.000 |

Population size $N=10^4$ , initial frequency $q_0=0.01$ , $q_{eq}$ : expected equilibrium frequency, $q_{obs}$ : observed frequency after $10^4$ generations. Results are average of 10 runs.

